# Prior photon measurement density function constructed in a standard channel space for functional near-infrared spectroscopy

**DOI:** 10.1101/2025.08.31.673398

**Authors:** Yang Zhao, Li-Jiang Wei, Xue-Ting Sun, Fa-Rui Liu, Lei Zhang, Chao-Zhe Zhu

## Abstract

Photon measurement density function (PMDF) determines the measurement locations and their associated sensitivity to the hemodynamic fluctuations of an fNIRS channel, which is essential for pre-experiment optode arrangement and post-experiment signal analysis/results interpretation in fNIRS studies. However, obtaining the exact PMDF relies on individual MR images, which are unavailable in common fNIRS studies. Previous studies alternatively adopt the PMDF derived from single head templates, i.e., template-PMDF, which ignores the anatomical difference between the template and the participants, as well as the anatomical variability across participants. Here, we propose a methodological framework using a standard channel space to construct a prior-PMDF from a structural MRI database for arbitrarily given channel locations. We demonstrate the utility of the prior-PMDF in estimating channel-wise fNIRS imaging parameters, such as subject-specific PMDF, sensitivity in intracerebral and extracerebral tissue compartments, and sensitivity to regions defined by brain atlas. The performance of the prior-PMDF tested on an independent dataset was compared with the template-PMDF. The results show that the prior-PMDF significantly surpasses the template-PMDF.

## 1. Introduction

Functional near-infrared spectroscopy (fNIRS) is a non-invasive neuroimaging technique that has been widely utilized in both scientific research and clinical situations in recent years. Compared with fMRI, fNIRS benefits from higher ecological validity, less sensitive to head motion, higher portability, and cost efficiency. It has higher spatial resolution than EEG and temporal resolution than fMRI. Due to these advantages, it has been prevalently applied in studies of special populations including infants and the elderly, as well as in studies requiring naturalistic experimental setups [Duan et al., 2013; Liu et al., 2021; Pinti et al., 2020; Quaresima and Ferrari, 2019].

fNIRS measures brain activation based on the complex interactions between light and the tissue of the human head. In the fNIRS experiment, a channel is often defined as the basic measurement unit, which includes a light source and a detector placed within a certain range of distance (often about 1~4 cm) on the scalp. The light source emits near-infrared light, which is received by the detector and forms the “banana-shaped” photon pathway, which covers spatially extended locations of tissues [Feng et al., 1995]. The photon density of a location in this pathway determines the sensitivities of the channel to the hemodynamic fluctuations at the location. The spatial distribution of the photon density in different locations is commonly termed the photon measurement density function (PMDF) [Arridge, 1995; Arridge, 1999; Arridge and Schweiger, 1995].

Since the PMDF of a channel determines its measured locations and their corresponding sensitivities, it has been incorporated in both pre- and post-experiment procedures of fNIRS studies. Based on the PMDF estimated on MR images of a participant, the sensitivity of a channel to the activation of the regions of interest (ROIs) can be estimated by summing the PMDF whose locations are within the ROI, which enables optimizing the channel arrangement on the scalp before an fNIRS experiment [Brigadoi et al., 2018; Machado et al., 2014; Morais et al., 2018; Wijeakumar et al., 2015]. The sensitivity to the extra-cerebral tissue, also termed the partial path length (PPL) in the tissue, is used to determine the optimal distance of short channel arrangement [Brigadoi and Cooper, 2015]. In the post-experiment data analysis, the sensitivity in the intra-cerebral tissues is the parameter used to convert the optical signal to hemodynamic concentration changes based on the modified Beer-Lambert law. The PMDF has also been used to improve the accuracy of activation analysis [Zhai et al., 2020]. Therefore, obtaining the PMDF is crucial for fNIRS studies.

The derivation of the PMDF involves photon propagation modeling based on an MR image, which commonly includes tissue segmentation and numerical methods such as the Monte Carlo photon simulation [Boas et al., 2002; Wang et al., 1995]. However, the individual MR image is commonly unavailable in fNIRS studies [Cutini et al., 2011]. Therefore, previous studies adopt acquiring and utilizing the PMDF based on the photon modeling on substitutional head templates, e.g., Colin27 or MNI152, namely the template-PMDF [Cooper et al., 2012; Custo et al., 2010; Morais et al., 2018]. However, using template-PMDF ignores the anatomical differences between the template’s head and the heads of participants recruited in an fNIRS study, as well as the anatomical variance across the heads of the participants. Using the template-PMDF can cause bias in predicting the fNIRS imaging parameters and further lead to suboptimal pre-experiment arrangement and post-experiment data analysis and result interpretation in an fNIRS study [Cooper et al., 2012; Custo et al., 2010]. Therefore, increasing PMDF estimation accuracy without using the individual MR image is desired for the fNIRS community.

Theoretically, the unbiased estimation of fNIRS imaging parameters can be achieved by directly averaging the parameters derived from multiple individuals. Therefore, in this study, we propose a methodological framework that enables synthesizing PMDFs of multiple subjects of an MRI dataset in a standard brain space, namely the prior-PMDF, given an arbitrary channel placement based on a previously proposed standard channel space. The channel space encourages characterizing arbitrary channel placement and corresponding same channel placement across different subjects [Wei et al., 2025]. This framework incorporates individual variability and potentially leads to more accurate estimations of important fNIRS imaging parameters. This paper is organized as follows: the methodological framework is first illustrated to construct the prior-PMDF based on PMDFs derived from MR images of multiple subjects. We then demonstrate the utilities of the prior-PMDF in estimating imaging parameters such as sensitivity in the brain and regions of a brain atlas. The prediction performance of the prior-PMDF for unseen subjects are also evaluated and the results are compared with that of the template-PMDF.

## 2. Theory and Methods

### 2.1. Theory of prior-PMDF construction

The methodological framework for constructing the prior-PMDF is shown in Figure 1. Since the individual MR images of an unseen subject are unavailable, the prior-PMDF should be constructed in a standard brain space, e.g., the MNI space, where probabilistic anatomical information is available. Therefore, the process of constructing the prior-PMDF is establishing a mapping from a specific channel placement on the scalp to its corresponding PMDFs in the standard space, i.e.,

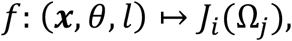

where the channel placement is characterized by the channel coordinate system, which includes three parameters: the channel location ***x*** (the middle point of a pair of source and detector), the orientation *θ* and the source-detector distance *l*. The channel location ***x*** is described by the continuous proportional coordinates (CPC): [*pNz, pAl*] [Xiao et al., 2018]. The orientation *θ* is the angle to the reference orientation (red arrow in Figure 1A) which is defined by the Scalp Geometry-based Parameter (SGP) [Jiang et al., 2022]. *J*_*i*_ (Ω_*j*_) is the sensitivity of a discretized region Ω_*j*_ (*j* = 1,2, …, *M*), commonly a voxel, of the *i*th subject in the MNI space. The channel coordinates of a specific fNIRS channel in the physical space of an individual can be obtained by manual measurement or using a 3D digitizer and a scalp navigation system. Based on the channel coordinates, the channel is transferred to the individual MR images in the MRI database to obtain the individual locations of source and detector, i.e., *s*_*i*_ and *d*_*i*_, where *i* is the *i*th individual in the database (Figure 1B). The individual photon fluence distributions, or the 2-point Green’s functions, of densely sampled scalp locations can be pre-computed based on the photon propagation modeling on individual MR images and registered to the MNI space based on the transformation of the anatomical registration [Custo et al., 2010; Yao et al., 2018]. In this way, the individual PMDFs in the database can be obtained rapidly without the time-consuming Monte Carlo simulation and the warping process. The individual PMDF can be derived using the adjoint method in the standard brain space, i.e.,

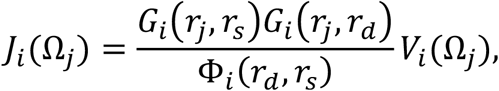

where *G*_*i*_(*r*_*j*_, *r*_*s*_) and *G*_*i*_(*r*_*j*_, *r*_*d*_) are the 2-point Green’s functions of the *i*th individual, at the location *r*_*j*_ produced by a source at the location *r*_*s*_ and a source at the location *r*_*d*_ respectively (Fig. 1D). Ω_*j*_ is a discretized region where *r*_*j*_ ∈ Ω_*j*_. Φ_*i*_(*r*_*d*_, *r*_*s*_) is the 2-point Green’s functions at the location of the detector produced by the source. *V*_*i*_(Ω_*j*_) is the volume of the voxel. Commonly used fNIRS parameters can be derived based on the individual PMDFs and anatomies. For example, the prior-PMDF for a channel placement can be derived in the common space as:

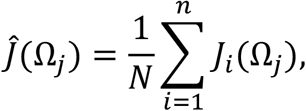

where *N* is the number of individuals involved in constructing the prior-PMDF. The statistics of sensitivity to regions of interest (ROIs) can be estimated by applying a mask of ROI in the standard space, i.e.,

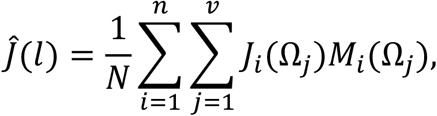

where *M*_*i*_(Ω_*j*_) is the mask of the ROIs in location Ω_*j*_ of the *i*th individual. The ROI could be a tissue compartment, e.x., the gray matter, a region defined by brain atlas, or a manually labeled area on the MNI brain. The masks can be the same across different individuals, as they are aligned to a common space, which implies *M*_*p*_(Ω_*j*_) = *M*_*q*_(Ω_*j*_) for ∀*p, q*. This procedure can be finished rapidly using MATLAB matrix multiplication.

**Figure 1.**
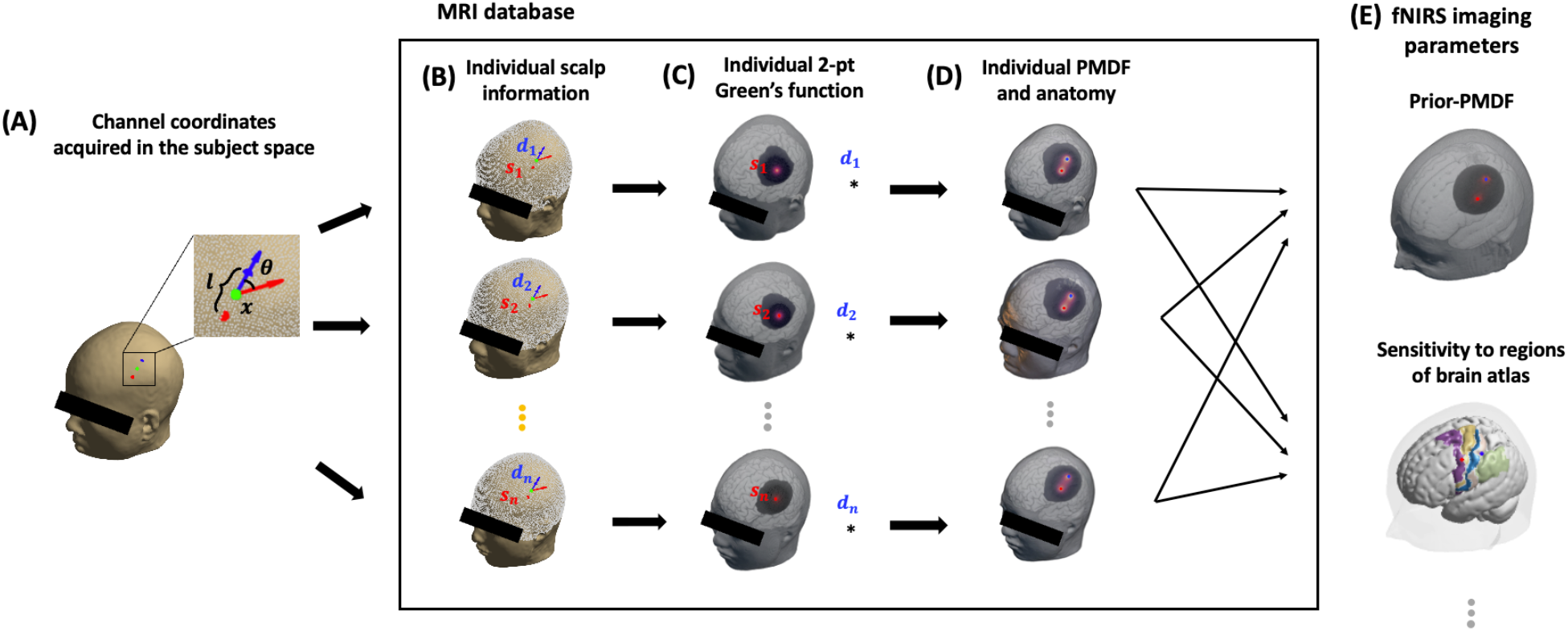
Flow chart of constructing the prior-PMDF given a specific channel placement. (A) A channel placement is described by the channel coordinate system, which includes three parameters: channel location ***x*** (green dot), orientation *θ* (blue arrow) and source-detector distance *l*. (B) Acquiring individual source and detector locations in the database based on the channel coordinates and individual scalp information. The white dots denote the high-resolution scalp locations. (C) Pre-computed Individual 2-point Green’s functions in a common brain space. (D) Individual anatomy and PMDF were derived using the adjoint method. (E) Predicted subject’s imaging parameters which include the prior-PMDF and sensitivity to regions defined by brain atlases.

### 2.2. Construction of prior-PMDF based on an MRI dataset

#### 2.2.1. Individual MR images and Monte Carlo simulation

45 MR images of healthy young adults were used in this study, which was randomly selected from a publicly available MRI dataset of adults [Wei et al., 2018]. These MR images were segmented using the SIMNIBS software into five tissue types, i.e., scalp, skull, CSF, gray matter, and white matter [Thielscher et al., 2015]. 18000 uniform distributed scalp locations were derived on the resultant head mesh. Monte Carlo simulation was used to obtain the 2-point Green’s function for each scalp location based on the Mesh-based Monte Carlo simulation software (MMC) [Fang, 2010]. The optical properties of each tissue were assigned as in Table 1, which is the mean optical properties across the typically used wavelength in fNIRS studies according to [Strangman et al., 2013]. The details of the MRI images and Monte Carlo simulation can be found in [Wei et al., 2024].

**Table 1.** Tissue optical properties for the Monte Carlo simulation.

| | $\mu_a(\text{mm}^{-1})$ | $\mu_s(\text{mm}^{-1})$ | $g$ | $n$ |
| --- | --- | --- | --- | --- |
| Scalp | 0.017275 | 0.72 | 0.01 | 1 |
| Skull | 0.011925 | 0.92 | 0.01 | 1 |
| CSF | 0.0025 | 0.01 | 0.01 | 1 |
| Gray matter | 0.0195 | 1.1 | 0.01 | 1 |
| White matter | 0.0169 | 1.35 | 0.01 | 1 |

#### 2.2.2. Prior-PMDFs construction in the standard brain space

The individual source and detector locations are derived for given channel coordinates by measuring the geodesic distance on the scalp meshes. The individual PMDFs in the MNI space are obtained by performing the adjoint method. Specifically, the individual 2-point Green’s functions are transformed to the common space based on the non-linear transformation derived by anatomical registration to the MNI head template. Since the voxel density is higher than the mesh density used in the Monte Carlo photon simulation, the mesh was unsampled using the iso2mesh toolbox therefore enabling one-to-one correspondence between the tetrahedrons and voxels. The two-point Greens’ functions on the mesh in the common space were directly mapped to the voxels based on the point-in-tetrahedron test. The volumes of mesh in the original space were accumulated to the voxels to preserve the subject-specific local tissue size. The normalization factor was also derived in the original space and saved. To reduce the cost of computation and storage, the prediction accuracy of different numbers of individual PMDFs involved in constructing the prior-PMDF was evaluated, which is detailed in the supplementary material (Section 6.1). The increase in the prediction accuracy is negligible when the dataset size is higher than 10. Therefore, a dataset size of 10 is used for constructing the prior-PMDF in the following analysis.

### 2.3. Prediction performance evaluations

To evaluate the prediction performance of the prior-PMDF framework, the estimated fNIRS parameters of typical probe placements were used as predictors and compared to that of the subjects in an independent dataset (n=10). The performance was evaluated for channels placed on typical scalp locations, i.e., the international 10/10 points, which include the following channel coordinates: for ***x***, 10/10 scalp locations, for *θ*, 0 and 90 degrees, for *l*, 30 mm (Fig. 2). The 10/10 locations on the rim were not included since some of their associate optode locations are unusual, such as on the ear or the eye. The parameters of the subjects in the independent dataset, i.e., the ground-truth value, were derived following the procedures in Section 2.2. First, the mean absolute error (*MAE*) of sensitivity to voxels is defined as:

**Figure 2.**
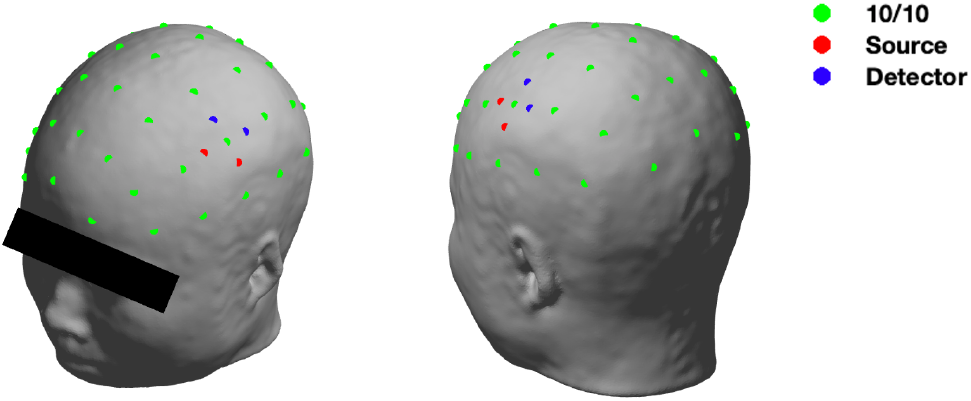
Scalp locations for evaluating the prior-PMDF. 10/10 scalp locations used for evaluating the prediction performance (Green points). Two exemplary channels are formed by two optode pairs with orthogonal orientations. Red and blue dots stand for the source and detector respectively.

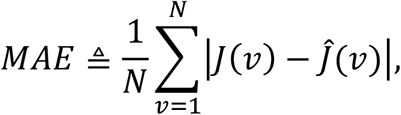

where *J*(*ν*) and 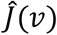 is the ground-truth and the predicted value of sensitivity to the voxel *ν* respectively. *N* is the number of voxels involved in the estimation. Note since the sensitivity value of one voxel is usually very small in the intracerebral areas, less than 0.1, the *MAE* would be near 0 when the evaluation involves the whole brain area. Therefore, only voxels whose ground truth value was larger than a certain threshold were involved when estimating the *MAE*. Second, the sensitivity to brain regions is often used to provide anatomical information on the functional activation and facilitate probe design [Brigadoi et al., 2018; Fu and Richards, 2021; Morais et al., 2018; Wijeakumar et al., 2015]. Therefore, the absolute error of the sensitivity to regions defined by a commonly used brain atlas, i.e., the Brodmann areas, is defined as:

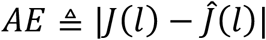

where *J*(*l*) and 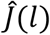 is the ground-truth and predicted sensitivity to the regions labeled as *l*. Since one channel can have sensitivities to multiple regions. The region with maximum sensitivity is used in the evaluation. Finally, since the sensitivity to intracerebral tissues is often utilized in converting optical signal to the hemodynamic concentration, i.e., the modified Beer-Lambert law [Ye et al., 2009]. The sensitivities to intra and extracerebral are also used in determining the distance of short channels and removing noises in extracerebral tissue compartments [Brigadoi and Cooper, 2015]. Therefore, the prediction error of sensitivities to intra and extracerebral are also evaluated.

The prediction performance was quantitatively compared with that derived by the template-PMDF. The PMDFs of two commonly used head templates, i.e., Colin27 and MNI152, are also derived for predicting subject-specific parameters. The same procedure of segmentation and photon propagation modeling as described in Section 2.2 was followed for the derivation of the template-PMDF. For the MNI152 atlas, the segmented MNI152 template in the Array Designer software was adopted [Brigadoi et al., 2018].

## 3. Results

### 3.1. Prediction of subject-specific PMDF

Figure 3. depicts the qualitative prediction performance of the prior-PMDF using a randomly selected example of a 30 mm channel placed on C3. The spatial pattern of the prior-PMDF is comparable with that of the individual PMDF (**Figure 3**A). The contour plot of the PMDF shows that the value of the PMDF decreases exponentially with the increase of the depths. **Figure 3**B shows the PMDFs resliced using the channel coordinates (green dash line in Fig. 4A). The spatial patterns of the individual PMDF and the prior-PMDF are comparable in different depths (**Figure 3**B). One can observe from the tissue template (third column) that the cerebral region mainly existed in the depths of 15 to 20 mm, in where the value of PMDFs is more matched than in other regions. The overall spatial pattern of the prior-PMDF is more dispersed than the individual PMDF.

**Figure 3.**
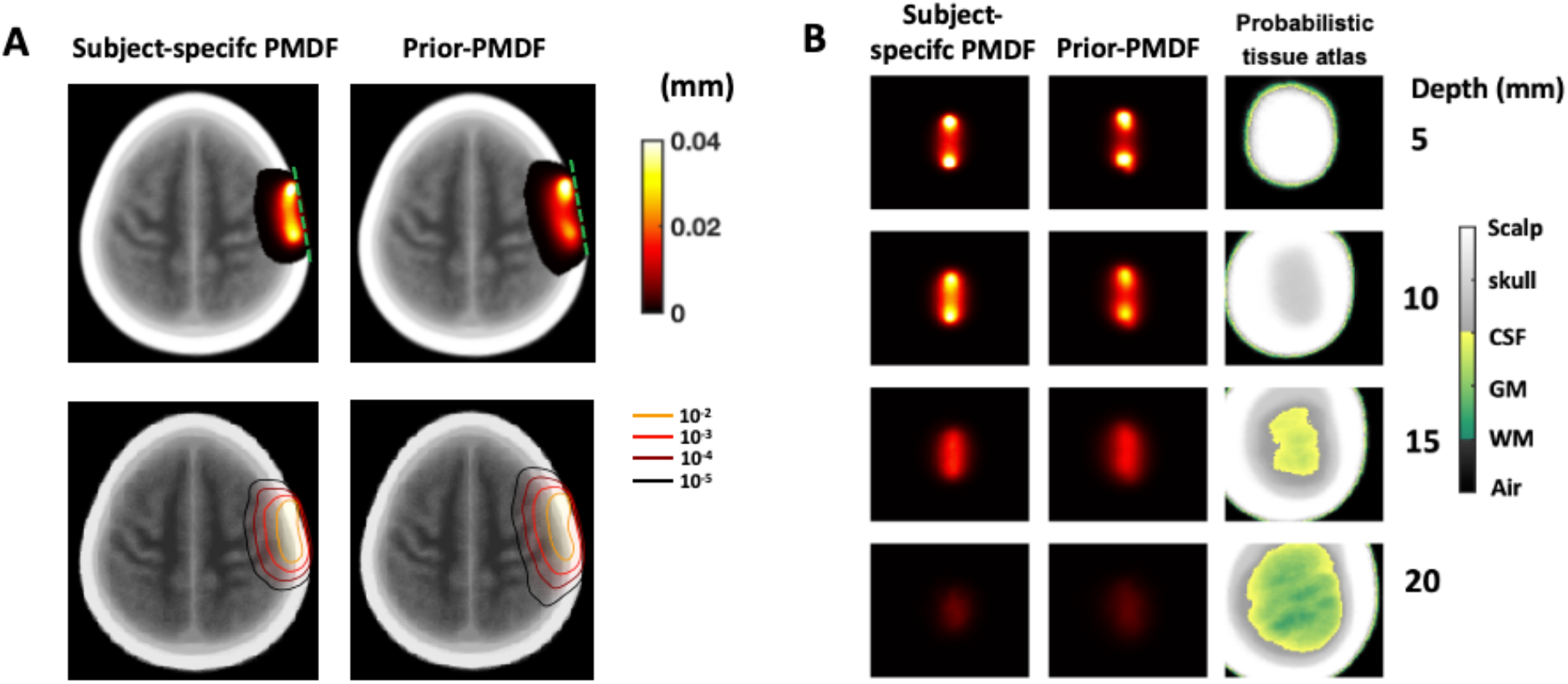
Prediction of subject-specific PMDF using the prior-PMDF. (A) A typical voxelized PMDF and a prior-PMDF with the same channel coordinates 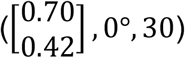. (B) PMDFs’ slices along the channel axis (green dash line in A) with different depths are shown with their associated anatomical information (last column).

### 3.2. Channel-sensitivity to brain regions

The predicted and subject-specific brain regions measured, and their corresponding sensitivities are shown in Figure 5. Regions with sensitivity greater than 5% of the total sensitivity (in intracerebral regions) were color-coded. The opacity indicates the percentage of the sensitivity to a region which is normalized to 1 by dividing the maximum sensitivity. Five regions, including the somatosensory motor cortex (BA 3, 1, 2), the primary motor cortex (BA 4), the premotor cortex (BA 6), and the supramarginal gyrus (BA 40), are predicted when the optodes are placed on C3, i.e., 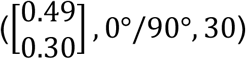 (Fig. 5, top row). The same regions are derived based on the subject-specific PMDF and their corresponding sensitivities are comparable (Fig. 5A, B, Fig. 5C, D). Interestingly, changing the orientation of the optodes changes the order of sensitivity to the measured regions, i.e., the sensitivity to the premotor cortex surpasses that to the primary motor cortex by placing the optodes paralleled to the Nz-Iz line. Only two regions, BA7 and angular gyrus (BA 9), are measured when optodes are placed on P1, i.e., 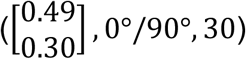 (Fig. 5, bottom row). The sensitivity to BA7 is much higher than that to angular gyrus and changing orientation slightly affects the sensitivity. This indicates that placing optodes on P1 has a better specificity to Brodmann areas compared with placing optodes on C3.

**Figure 5.**
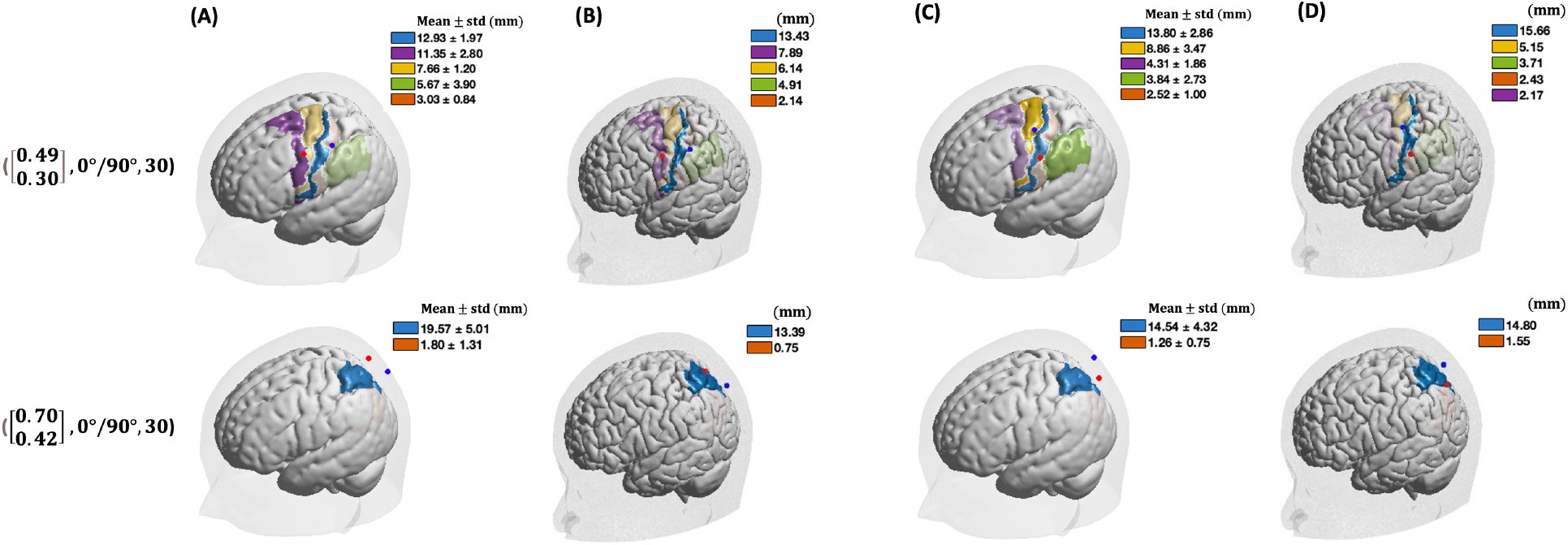
Sensitivity to the Brodnman areas for optode placement at channel coordinates. 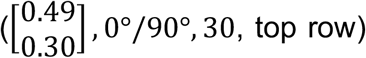 **and** 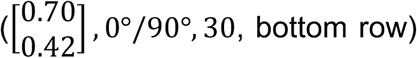. Both the predicted (A and C) and a randomly selected subject-specific (B and D) sensitivity are depicted. Color-coded regions indicate sensitivity > 5%. Blue and red dots represent the source and detector respectively.

### 3.3. Prediction of sensitivity to intra- and extracerebral regions

The value and spatial distribution of the predicted sensitivity to intra- and extracerebral tissues using the prior-PMDF are depicted with that derived by the subject-specific PMDF (Fig. 6). The spatial distribution shows the sensitivity to the intracerebral tissues is higher in the frontal lobes compared with the parietal lobes, which is in accordance with literature studies [Strangman et al., 2014]. The spatial pattern of the sensitivity is comparable between using the prior-PMDF and a randomly selected subject-specific PMDF. The spatial pattern of the sensitivity derived by the prior-PMDF is also smoother than that derived by the subject-specific PMDF, which is mainly caused by the averaging process.

**Figure 6.**
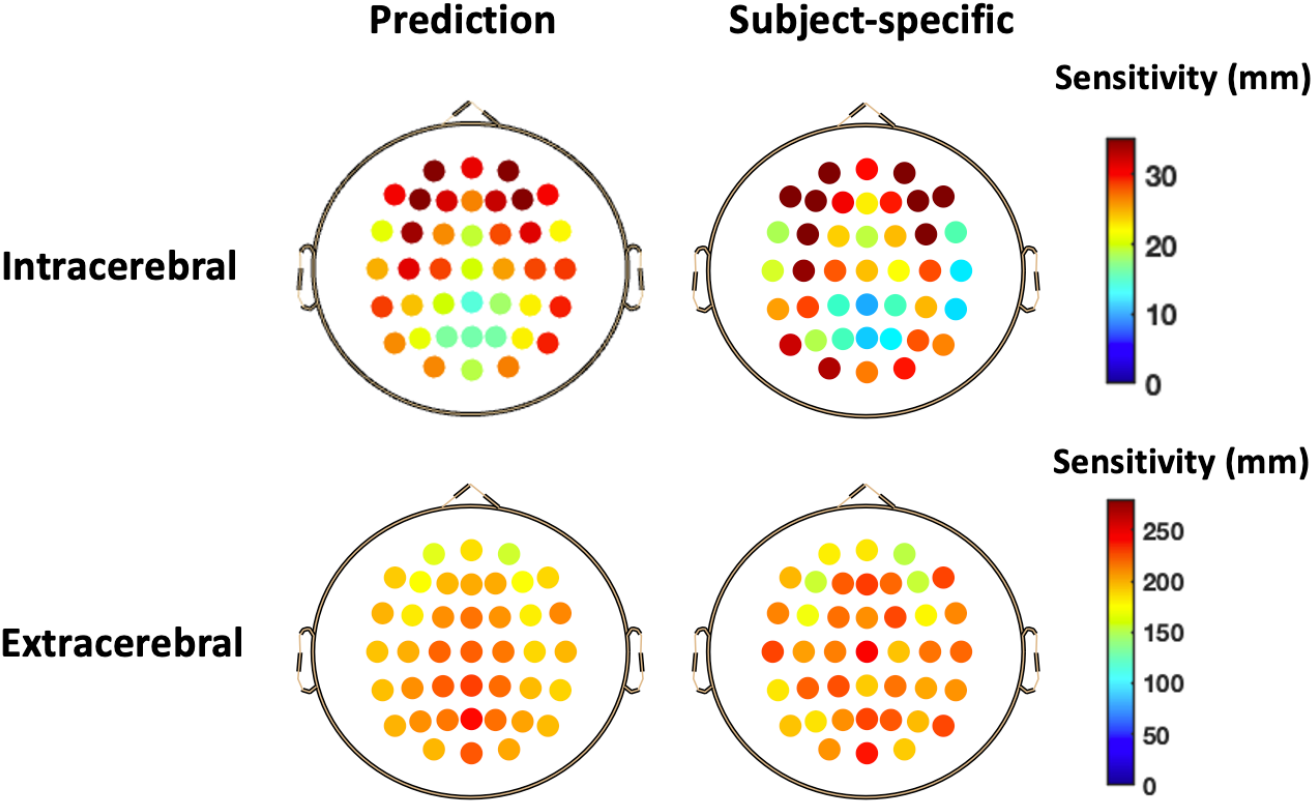
Prediction of sensitivity to the intra- and extracerebral tissues using the prior-PMDF. The sensitivity to intracerebral estimated on the 10/10 scalp locations using the prior-PMDF (left) is compared to that using a randomly selected subject-specific PMDF (right).

### 3.4. Prediction performance comparison with template-PMDF

The prediction errors of the parameters using the prior-PMDF are quantitatively compared to that of the template-PMDF (Fig. 7). The absolute error of the sensitivity to the Brodmann areas predicted by the prior-PMDF is 4.64 ± 0.75 mm (mean ± std), which is significantly lower than that predicted using the PMDFs of Colin27 (8.38 ± 1.90 mm) and MNI152 (11.07 ± 2.03 mm; P<0.001). The absolute error of the sensitivity to intracerebral tissues predicted by the prior-PMDF is significantly lower than that predicted by the template-PMDF (5.27 ± 1.03 mm vs 16.89 ± 3.58, Colin27, 11.75 ± 3.17 mm, MNI152;

**Figure 7.**
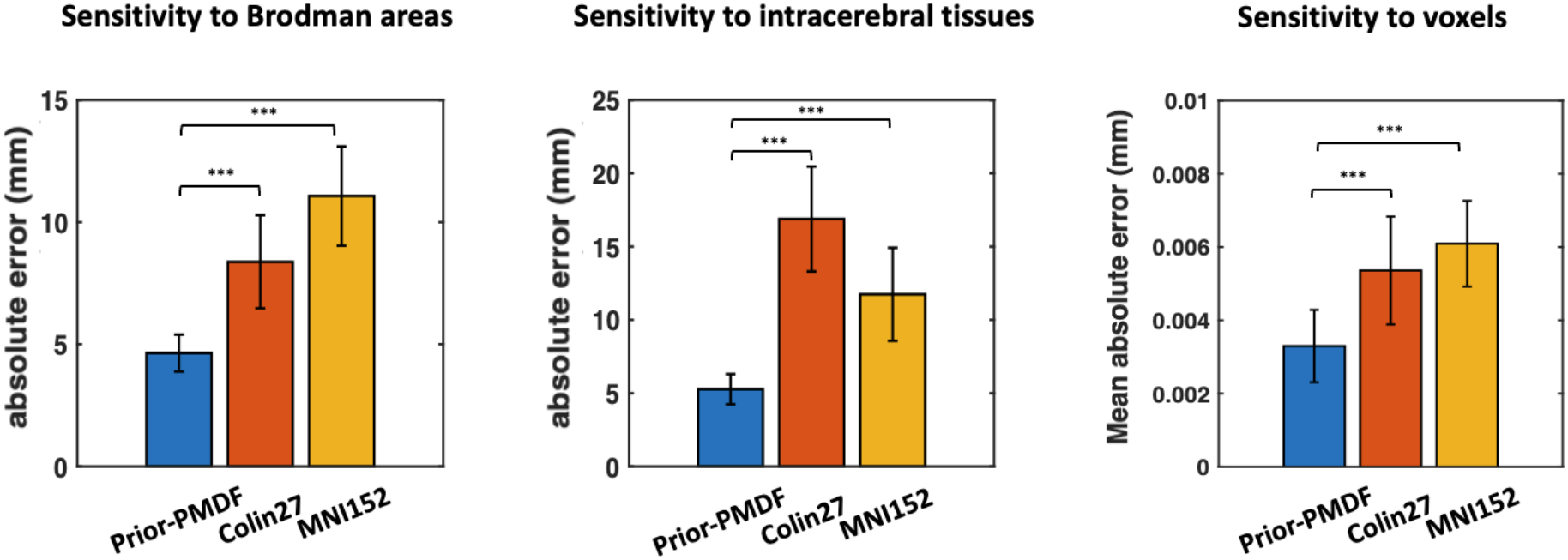
Quantitative comparison of the prediction error of the parameters using the prior-PMDF to that using the template-PMDF. The Sensitivities to the Brodman areas (left), intracerebral tissues (middle), and voxels (right) are compared to those of the template-PMDF. The error bar stands for the standard deviation across subjects. *** indicates p<0.001 in a two-tailed paired *t*-test (N=10).

P<0.001). For the sensitivity to voxels, the evaluation locations include gray matter voxels (determined by the individual gray matter mask) whose ground-truth values are larger than 0.01, which makes a primary contribution to the fNIRS signal. The absolute error of the prior-PMDF is 0.33 ± 0.10 mm (mean ± std), which is significantly smaller than that of the Colin27 ([0.54 ± 0.15] *10^-2^ mm; P<0.001) and the MNI152 ([0.61±0.12] *10^-2^ mm; P<0.001). The standard errors of the prediction errors of the prior-PMDF are also smaller than those of the template-PMDFs.

Since the prior-PMDF supports arbitrary channel arrangements, the prediction error of the prior-PMDF was further evaluated and compared with that of the template-PMDF across different source-detector distances (Table 2). Only distances longer than 15 mm are shown as anything shorter would lead to very low sensitivity to the intracerebral regions. The results show that the prior-PMDF has a lower mean absolute error in all the evaluated distances.

**Table 2.** Prediction error of sensitivity to voxels when channels are arranged with varied distances using the prior-PMDF and the template-PMDF.

| Distance | Mean absolute error ($\times 10^{-2}$ mm) | | |
| --- | --- | --- | --- |
|  | Prior-PMDF | Colin | MNI152 |
| 15mm | 0.58 | 0.84 | 0.78 |
| 20mm | 0.46 | 0.73 | 0.70 |
| 25mm | 0.40 | 0.64 | 0.64 |
| 30mm | 0.33 | 0.54 | 0.61 |
| 35mm | 0.29 | 0.47 | 0.62 |
| 40mm | 0.30 | 0.44 | 0.65 |
| 45mm | 0.31 | 0.44 | 0.68 |

## 4. Discussion

In this study, we propose a methodological framework that enables estimating fNIRS imaging parameters based on synthesizing PMDFs from multiple subjects in a database, the prior-PMDF. We demonstrated estimating subject-specific PMDF, sensitivity to Brodmann areas, and sensitivity to intra- and extracerebral tissues based on the prior-PMDF framework. The parameters predicted by the prior-PMDF are compared to those predicted by the template-PMDF. The results convergently showed the prediction performances of the prior-PMDF are higher than that of the template-PMDF.

One of the fundamental difficulties of the fNIRS technology is the unavailability of subject-specific anatomical information [Strangman et al., 2002]. Therefore, sophisticated photon migration modeling cannot be applied to acquire important imaging parameters to facilitate pre-experimental optode arrangement and post-experimental data analysis. In the fNIRS literature, methods have been proposed to localize the brain areas detected by the placed optode montages. Homan and Okamoto explored the scalp-to-brain correspondences in EEG scalp locations using a projection method, i.e., projecting the channel locations (defined in the middle between the source and the detector) to the brain surface [Homan et al., 1987; Okamoto et al., 2004]. Approaches such as the probabilistic registration and the transcranial brain atlas have been proposed later to enable localization of arbitrary scalp locations [Singh et al., 2005; Xiao et al., 2018]. These methods transcend the understanding of fNIRS results from the activation pattern on the scalp space to the brain space. However, they are criticized for using an approximated model, i.e., from a scalp point to a cortical point, to obtain the scalp-brain correspondence, which ignores the realistic principle of the fNIRS imaging in recent studies [Brigadoi et al., 2018; Cai et al., 2021]. The actual measurements of an fNIRS channel are reflected by the PMDF, whose pattern on gray matter exhibits non-uniformity across an extensive spatial domain. The PMDF patterns also varied across channels placed on different scalp locations. The point model cannot reveal these important informations, which can lead to suboptimal optode arrangements or misunderstanding of the fNIRS results. The fNIRS community is moving forward to acquire the PMDF given channel placements. Researchers can obtain the PMDF of channels based on the head template and design an optode montage for an ROI. However, as we show in this study, the template-PMDF has a lower prediction performance without considering the anatomical variability between the template and the participants, as well as the anatomical variability across participants.

The increased prediction performance of the prior-PMDF can be attributed to the consideration of the anatomical variabilities. First, compared with using a single template, using the averaged imaging parameters of multiple individuals reduces the prediction bias of the targeted participants in an fNIRS study as shown in Figure 3. The variance of head size and shape is incorporated by transferring the channel to multiple individuals in the MRI database using the channel coordinates. The difference in PMDF across participants due to the variance of tissue distribution is also incorporated. One confounding factor is that the templates are Colin27 and MNI152 which are mainly Caucasians, and the targeted participants are Chinese young adults which are Asians. We conducted another analysis to address this issue. Specifically, PMDFs were derived on a template constructed using the MR images in the current dataset and its prediction performance is compared to that of the prior-PMDF (Section 6.2). The results showed that the prior-PMDF still has significantly higher prediction accuracy than the template-PMDF, which indicates the increased performance is not due to the difference in population but to incorporate the anatomical variability across subjects in the dataset (Fig. S2). The prediction performance also varied across different scalp locations. The spatial map of the prediction error depicts a lower value on the frontal lobe compared with the parietal and occipital regions (Fig. S3). This is in line with the variability of the point-to-point scalp-brain correspondence across individuals found in the TBA, which is caused by lower anatomical variability in the frontal area than in the parietal and occipital areas.

There are two additional contributions from the current study. First, the volumetric PMDFs are provided for arbitrary channel placement, which supports broader applications than providing PMDFs on specific tissues, e.g., gray matter surface. For instance, the PMDF of extra-cerebral regions can be used to decide optimal short-channel separations. The sensitivity to the intra-brain region is used to convert the optical signal to the hemodynamic signal. Second, we provide a method to directly compare the PMDFs of different individuals in a common space, which can be used to compare the PMDFs of different populations, e.g., male versus female, adult versus the elderly.

Two limitations remain in the proposed method. First, we used the mean optical properties of multiple commonly used wavelengths. Future studies should enable obtaining prior-PMDF of different wavelengths, which can be achieved by evaluating the differences in PMDF affected by the wavelength. Second, the storage required for the proposed method is demanding since the individual 2-point Green’s functions of dense scalp locations are necessary to be stored. Methods such as sparse representation or principal component analysis can be incorporated to reduce the storage burden required for applying the current method.

In future studies, the prior-PMDF can be applied and evaluated in more applications both in pre- and post-experiment procedures. For example, the sensitivity to the intracerebral tissue is different across different scalp locations. The optode can be arranged with different separations, according to the prior-PMDF, to encourage measurement homogeneity across different scalp locations. The predicted sensitivity can also be used as the parameter for signal conversion in the post-experiment analysis to promote quantification of the hemoglobin concentration change. The performance of using prior-PMDF in the image reconstruction of DOT can be evaluated and compared to that of using the template-PMDF. Finally, since the PMDF can be varied across different populations, such as populations with different ages or genders, the age-depend prior-PMDF can be constructed for specific populations, which can facilitate fNIRS studies targeted to specific populations and enable the consideration of confounding factors in the studies focusing on comparing brain functions between different populations.

## 5. Conclusion

We proposed a methodological framework to enable constructing the prior-PMDF for arbitrary scalp channel placement, which incorporates inter-subject anatomical variability.

The prediction of fNIRS imaging parameters including the subject-specific PMDF, sensitivity to brain regions, and sensitivity to intra- and extracerebral tissues were demonstrated. The prior-PMDF has significantly higher prediction performance in predicting these imaging parameters than the template-PMDF. Being able to provide more reliable and accurate fNIRS imaging parameters, the prior-PMDF can improve the quality of pre-experiment optode arrangement and post-experiment signal analysis in future fNIRS studies.

## Author contributions

**Yang Zhao**: Conceptualization, Methodology, Data Curation, Formal analysis, Writing - Original Draft. **Li-Jiang Wei**: Methodology, Data Curation. **Xue-Ting Sun**: Data Curation, Resources. **Fa-Rui Liu**: Data Curation. **Lei Zhang**: Conceptualization, Supervision, Review & Editing. **Chao-Zhe Zhu**: Conceptualization, Supervision, Review & Editing.

## Data and code availability statements

The data and code that support the findings in this study can be available by contacting the corresponding authors, Prof. Chao-Zhe Zhu or Prof. Lei Zhang, upon a formal data sharing agreement.

## Ethics statement

This study used publicly available data from the cross-sectional Southwest University adult lifespan dataset at 10.1038/sdata.2018.134, which does not contain any identifiable personal information and does not require additional ethical approval.

## Acknowledgment

This work was sponsored by the National Natural Science Foundation of China (82071999 and 61431002).

