## Supplementary materials for "Prior photon measurement density function constructed in a standard channel space for functional near-infrared spectroscopy"

### 6.Supplementary Materials

#### 6.1. Dataset size for constructing prior-PMDF

The number of individuals used in constructing the prior-PMDF is an important parameter, which may cause different prediction accuracy. Ideally, the prediction accuracy would increase with the increase of the dataset size involved in the construction. However, 2-point Green’s functions of dense scalp samples need to be stored for each individual in the hard disk, which increases the storage burden. Therefore, the different number of individual PMDFs for constructing the prior-PMDF was evaluated to determine the optimal dataset size. Specifically, 10 out of 45 individuals are randomly selected as a test dataset, in which the individual PMDFs are set as the golden standard. Other PMDFs of the left 35 individuals are selected in increments of 5, ranging from 1 to 35, and averaged to construct the prior-PMDF. The constructed prior-PMDF are then used to predict the ground truth PMDFs in the test dataset. The Pearson correlation coefficient is used to measure the similarity of the spatial pattern of PMDF predicted by the predicted PMDF and that of the ground truth, i.e.,

$$r=\frac{cov(J,\hat{J})}{\sigma_{J}\sigma_{\hat{J}}},$$

where $cov(J,\hat{J})$ is the covariance of the ground truth PMDF and predicted PMDF, $\sigma_{J}$ and $\sigma_{\hat{J}}$ is the standard deviation of the ground truth PMDF and predicted PMDF. The correlation coefficient is evaluated to determine the optimal dataset size. The prediction accuracy is evaluated on the channel coordinates described in Section 2.3 in this analysis (Fig. 2).

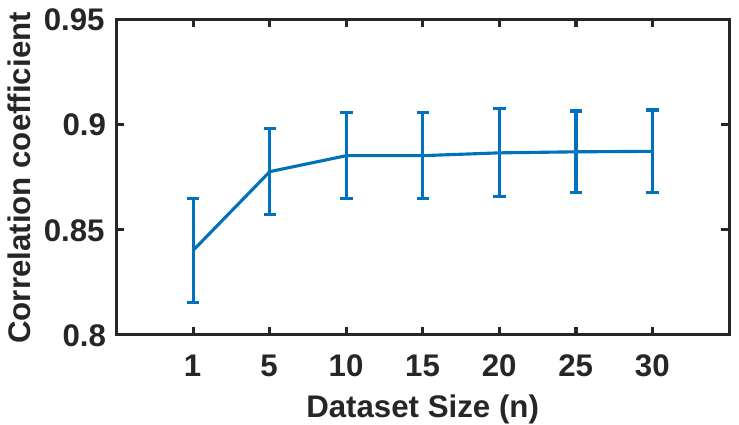

**Figure S1. Prediction accuracy of prior-PMDF constructed with different numbers of dataset size.** The error bar represents the standard error across 10/10 scalp locations.

The correlation coefficient increases with the increase in dataset size, which indicating the similarity of patterns between the predicted PMDF and ground truth PMDF increases (Fig. S1). The increase of correlation is negligible after the dataset size of 10. Therefore, a dataset size of 10 is used for constructing the prior-PMDF.

#### 6.2. Prediction performance comparison with Ch35-PMDF

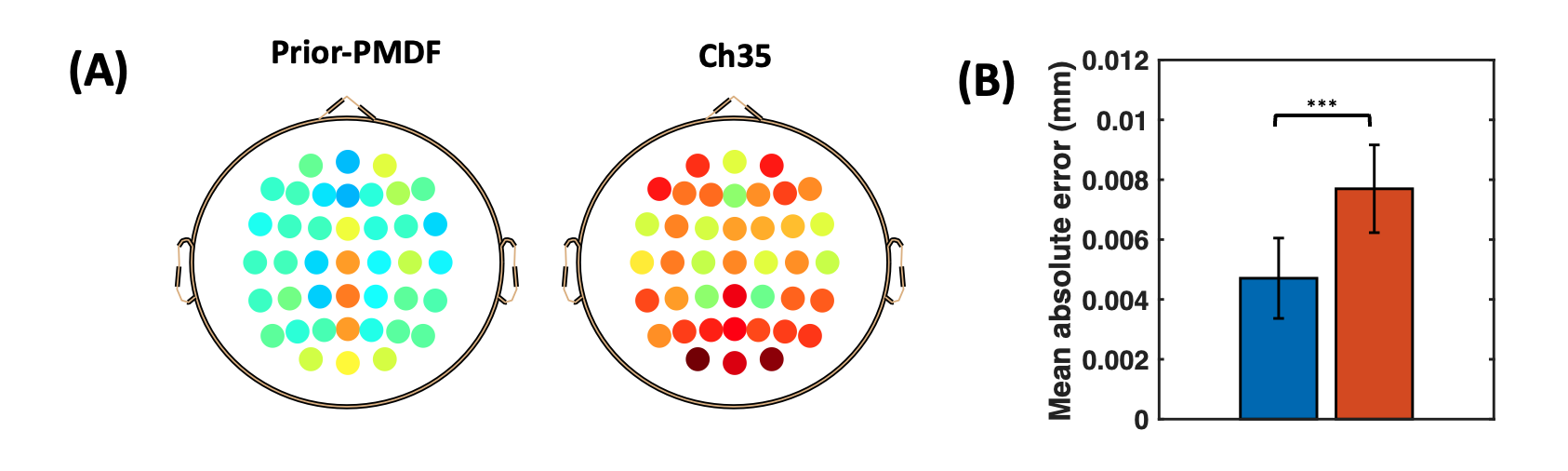

Figure S2. Prediction error of the prior-PMDF and the Ch35-PMDF. (A) Mean absolute error (MAE) across scalp 10/10 locations. (B) Statistical comparison of MAE between prior-PMDF and Ch35-PMDF. Error bars represent the standard deviations across the 10/10 scalp locations.

To compare the prior-PMDF with the template-PMDF, Monte Carlo simulation is performed on a head-template built using the structural MR images (N=35) in the same MRI dataset, termed the Ch35. The head template is constructed according to (ref) using the ANTS software. Specifically, the initial template is created by transforming individual MR images to the ICBM152 template using rigid transformation and averaging them across subjects. Then same individual MR images are transformed onto the initial template by nonlinear transformation and averaged across subjects. To obtain the segmented template, the individual segmented tissue images were transformed to the Ch35 template based on the individual nonlinear transformation. The tissue label of each voxel is determined as the most frequent tissue label across subjects. Then the 5 layer tissue meshes are derived by the make_surface_mesh and make_volume_mesh modules of the headreco module which is provided by the SIMNIBS software. Mesh-based Monte Carlo simulation (MMC) is run with same parameters as in Prior-PMDF construction on the resultant head mesh. The MAE of the prior-PMDF is lower and more consistent across scalp locations than that of the Ch35-PMDF (Fig. S2, A). The statistical results show the MAE of the prior-PMDF are significantly smaller than that of the Ch35-PMDF (paired t-test, p<0.001, Fig. S2, B).

#### 6.3. Prediction performance of prior-PMDF and template-PMDF

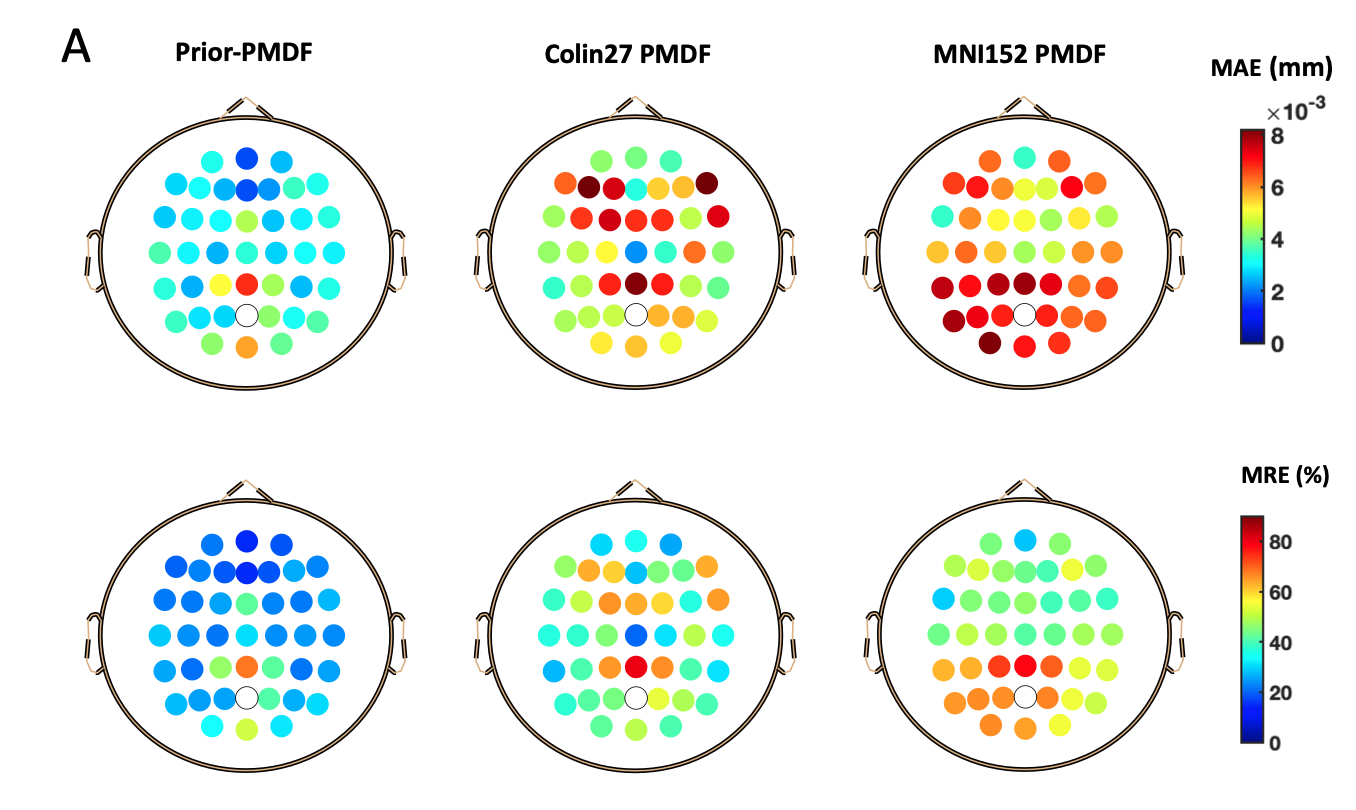

Figure S3. Prediction error of the prior-PMDF and the template-PMDF across scalp 10/10 locations.
